# The effects of varying intensities of unilateral handgrip fatigue on bilateral movement

**DOI:** 10.1101/2024.12.01.626257

**Authors:** Adrian L. Knorz, Justin W. Andrushko, Sebastian Sporn, Charlotte J. Stagg, Catharina Zich

**Affiliations:** Wellcome Centre for Integrative Neuroimaging, FMRIB, Nuffield Department of Clinical Neurosciences, University of Oxford, United Kingdom; Department of Sport, Exercise and Rehabilitation, Faculty of Health and Life Sciences, Northumbria University, Newcastle upon Tyne, United Kingdom; Department of Clinical and Movement Neuroscience, UCL Queen Square Institute of Neurology, London, United Kingdom; Medical Research Council Brain Network Dynamics Unit, Nuffield Department of Clinical Neuroscience, University of Oxford, United Kingdom

## Abstract

The human ability to maintain adequate movement quality despite muscle fatigue is of critical importance to master physically demanding activities of daily life and for retaining independence following motor impairments. Many real-life situations call for asymmetrical activation of extremity muscles leading to unilateral manifestations of muscle fatigue. Repeated unilateral handgrip contractions at submaximal force have been shown to be associated with neural dynamics in both contralateral and ipsilateral cortical motor areas and improved response times of the contralateral, unfatigued homologue in a button-press task. However, it remains unclear whether the observed improvement in contralateral response latency translates into higher-level benefits in movement quality.

To investigate this, 30 healthy participants underwent unilateral handgrip fatiguing tasks at 5%, 50%, and 75% of maximum voluntary contraction (MVC) force. Subsequently, bimanual movement quality was assessed in an object-hit task using a Kinarm robot.

The protocol at 50% and 75% of MVC elicited clear signs of muscle fatigue compared to the control condition (5%) measured by a decline in force, post-exercise deterioration in MVC, characteristic changes in surface electromyography magnitudes, and increases in ratings of perceived exertion. No change was observed on kinematic measures in the object-hit task for both arms indicating that unilateral handgrip fatigue did not elicit measurable effects on higher-level movement quality on the ipsilateral or contralateral homologue. Previously reported improvements on contralateral response latency were not found to translate into advanced movement quality benefits.

## 1. Introduction

The human body’s inherent adaptability in the face of muscle fatigue is vitally important for a wide range of daily activities. In healthy individuals, the ability to make effective adjustments allowing accurate movements to continue even during muscle fatigue is highly advantageous for a range of activities, including athletic performance and professions that require strenuous manual labour. Further, in patients with motor impairments, maintaining quality movements despite increased fatiguability is critical for retaining independence in daily activities and avoiding the need for external care.

Muscle fatigue is a complex and often rather vaguely defined physical state, which can arise through a number of peripheral and central processes. While it can sometimes be broadly defined as a ‘transient decrease in the capacity to perform physical actions’^1^ it is often characterised in a more focused way as an ‘exercised-induced decrease in the ability to produce force’.^2^ Muscle fatigue can be quantified by various physiological markers, among them a decline in force output and distinctive electromyographical alterations.^1^ Muscle fatigue generally develops gradually, and fatigue-induced changes in physiological markers are commonly at least partly task-dependent.

While muscle fatigue can occur symmetrically, many real-life situations lack perfect symmetry, meaning that even healthy individuals regularly face asymmetric stresses on their motor system. For example, on a crowded bus, a person might choose to tightly grip a passenger handle with one hand for the entire duration of the bus ride. After descending the bus, the same individual might then choose to perform a more complex movement using their contralateral, unfatigued arm (e.g., inserting a key into the door of their house) or even engage in bimanual compound movements of both arms (e.g., taking off their coat and untying their shoes). To predict how the individual’s preceding unimanual fatigue will now impact the movement quality of such contralateral or bilateral movements is complex and involves changes in underlying neural dynamics which might have been triggered by the unimanual fatigue, and have the potential to not only influence the fatigued limb itself but also the contralateral, unfatigued homologue.

Recent evidence from our research group suggests that repeated unilateral handgrip contractions at 50% of maximum voluntary contraction (MVC), resulting in a decrease in volitional force output, are associated with an improvement in motor response latency in the contralateral, unfatigued homologous limb with a button-press task compared to a control condition of 5% of MVC.3 Further, functional magnetic resonance imaging (fMRI) and magnetic resonance spectroscopic imaging (MRSI) revealed that the degree of change in gamma-aminobutyric acid (GABA) in the ipsilateral primary motor cortex (M1) and the bilateral supplementary motor areas (SMA) after the contractions at 50% of MVC were in relation to the contralateral response time improvement.3

However, it is still incompletely understood which range of fatigue intensities (i.e. percentages of MVC) can trigger such neural responses, given that previous experiments have been conducted using only one fatigue intensity (e.g., 50% of MVC vs. 5% of MVC as a control). Furthermore, even though we have shown that the degree of decline in GABA following unilateral handgrip contractions is related to improvements in contralateral limb response time, it is unclear whether this effect translates into quantifiable benefits in higher-level movement quality, as opposed to only in response time. This is particularly relevant when considering movement accuracy. While the disinhibitory effects of decreased GABA may benefit performance in terms of reducing movement response time, it is less certain whether this effect will also translate to benefit finer motor skills, which may not only require speed but also accuracy. Indeed, Branscheidt et al. have suggested that muscle fatigue may even be responsible for a longer-term effect of ‘target overshoot’ as a result of excessive force production^4^. In light of the principles of disinhibition, one might reason that these seemingly contradictory movement effects of muscle fatigue could in fact represent two sides of the same coin. Downregulation of GABA could theoretically account for both effects, which would create an interesting trade-off: beneficial effects on response time and overall force production at the expense of movement accuracy and possible target overshoot.

Previous research has also indicated that there are potentially conflicting effects transpiring in the supraspinal realm as a response to unilateral muscle fatigue. Previous protocols of unilateral fatigue in the right dorsal interosseus muscle^5^ and the knee extensors^6^ have been reported to induce various degrees of cross-over effects of increasing fatigue on the contralateral homologue, while a protocol of right unimanual fatigue found no evidence for a cross-over effect.^7^ A TMS study of unilateral handgrip contraction exercise on the left hand reported decreased short interval intracortical inhibition (SICI) in the primary motor cortex (M1) on the ipsilateral side and suggested diminished excitability of the ipsilateral M1^8^, contrasted by other results suggesting that forceful left abductor pollicis brevis (APB) exercise at intensities of more than 50% of MVC might be associated with anti-inhibitory effects on the contralateral homologue.^9^

One probable reason for the inconsistent evidence is the heterogenous nature of employed experimental protocols.

When considering real-world implications, it is also difficult to infer predictions about the movement quality of bilateral compound movements from purely unilateral measurements. As Swinnen^10^ has accurately noted in 2002, ‘principles of interlimb coordination are unique and cannot be inferred from the laws of single-limb movements’. Feeney et al^11^ meanwhile noted the importance of those coordinated interlimb movements for the fatigued state, reasoning that ‘switching to bimanual tasks when one hand becomes fatigued could be beneficial regarding preserving the high level of both the manipulation performance and force coordination’.

Therefore, in this study we aimed to elicit varying intensities of unilateral handgrip fatigue and measure how these influence movement quality as assessed by advanced kinematic parameters of both arms during bilateral movements. We deliberately designed our experiments to require not only speed but also fine motor accuracy, creating experimental leverage in identifying problems of overshoot and rapid bimanual coordination.

## 2. Methods

### 2.1. Participants

We recruited 30 right-handed participants, between 19 and 41 years of age (median: 25, IQR: 10.75), of which 18 (60%) were female. Informed consent was acquired in written form from all participants. This study has been approved by the Oxford Central University Research Ethics Committee (CUREC) (R88022/RE003) and was performed in accordance with the Declaration of Helsinki.

### 2.2. Session overview

#### Acclimatisation and pre-assessment

Each participant completed a single session (∼ 2.5 hours duration). After informed consent, participants were first allowed to become accustomed to the KINARM robot. The chair height was adjusted to enable comfortable operation of the robotic handles, while the position of the chair was kept strictly centred. Participants underwent 1 min of training to familiarise themselves with the Assessment Task (AT).

To individualise the force levels for the Fatiguing Task (FT), each participant’s maximum voluntary contraction (MVC) was quantified. Participants performed 3 cued contractions at MVC, each of 3 sec duration with 30 sec of rest in between contractions (MVC 3×). Verbal encouragement was given during MVC attempts to facilitate maximum efforts and force output. By employing the 2×2 Achievement Goals Questionnaire for Sport (AGQ-S) by Elliot and McGregor^12^ directly before the MVC calibration, we aimed to further encourage participants and to quantify their performance motivation.

After quantifying individual MVC (via MVC 3×), participants completed a number of questionnaires: Edinburgh Handedness Inventory (EHI)^13^, World Health Organization Global Physical Activity Questionnaire (GPAQ)^14^ and BORG-CR-10^15^. Next, a single MVC (i.e., MVC 1×) was obtained followed by the baseline AT.

#### Main experiment

During the main experiment, participants performed three rounds of tasks, each round comprised the following: MVC 3×, Fatiguing Task (FT), MVC 1x, BORG-CR10, Assessment Task (AT), and a recovery period (**Figure 1a**). The only difference between the three rounds was the level of force intensity (5%, 50% or 75% of MVC) during the FT.

**Figure 1:**
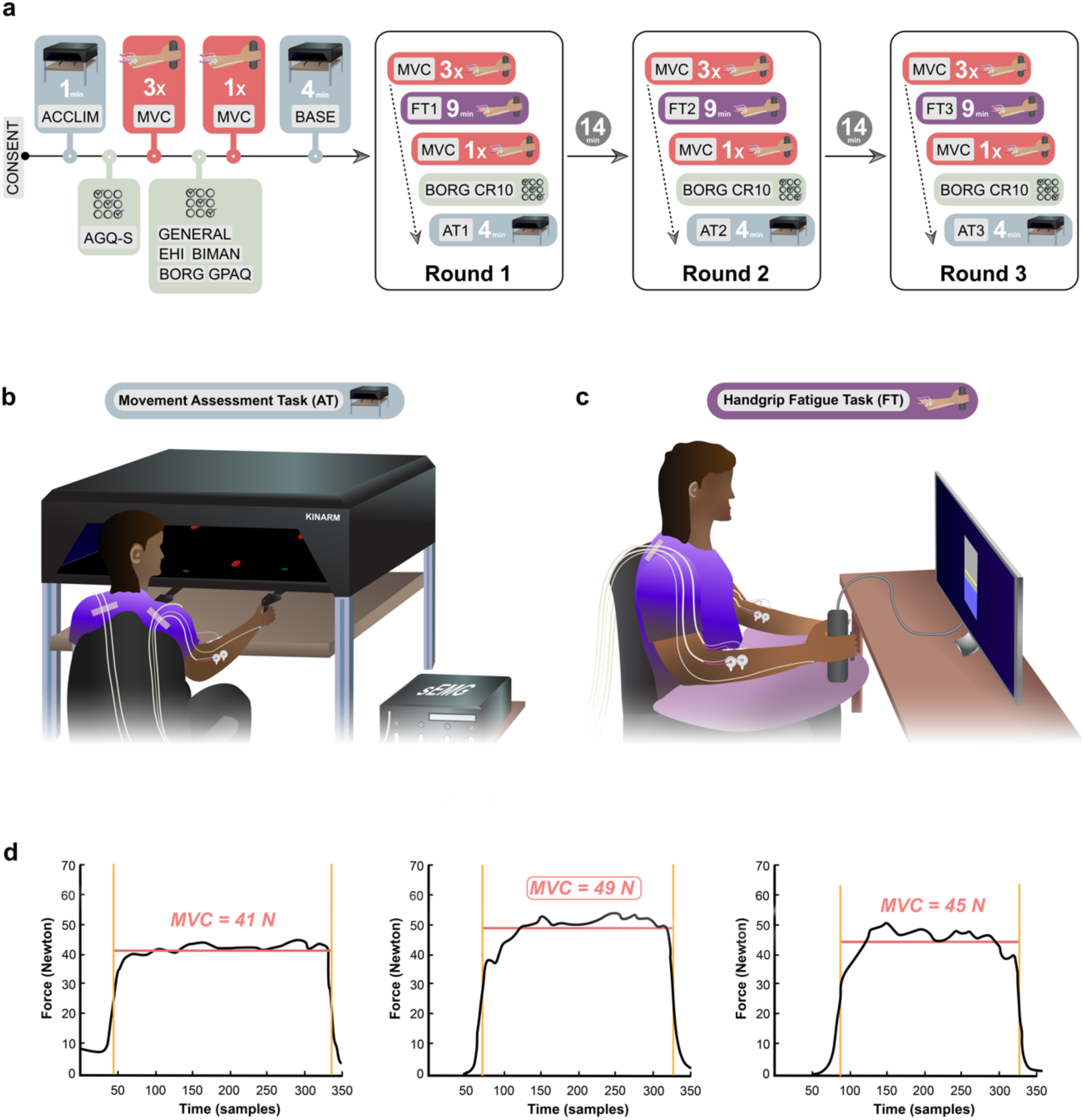
Experimental Design. a) Experimental timeline. Abbreviations: CONSENT=Informed consent of participants, ACCLIM=Kinarm acclimatization period, AGQ-S=Achievement Goals Questionnaire for Sport, MVC=Maximum Voluntary Contraction, GENERAL=General and Demographic Questionnaire, EHI=Edinburgh Handedness Inventory, BIMAN= Bimanual Tasks in Daily Life Questionnaire, BORG= Borg-CR10 rating scale, GPAQ=WHO Global Physical Activity Questionnaire, BASE=Kinarm baseline period. b) Experimental setup of the movement assessment task (AT). c) Experimental setup of the handgrip fatigue task (FT). d) Example of calculation of MVC 3×

#### Fatiguing task (FT)

We employed a visually prompted handgrip FT of 9 min duration, consisting of 1 sec handgrip contractions followed by 1 sec relaxation periods for a total of 270 contractions. Participants were instructed to adjust their applied handgrip force to match a target line displayed on a computer screen (**Figure 1c**). Participants were shown their handgrip force output in real time. The horizontal target line was kept consistently in the centre of the screen to maintain a consistent visual task presentation across the three conditions, regardless of the force intensity required (i.e., force intensity of 5%, 50%, or 75% of baseline MVC. Six protocols were created to balance the order of the three experimental conditions across participants. Participants were allocated to a protocol using covariate-adaptive allocation, employing a variance minimisation procedure developed by Sella et al^16^. This allowed us to assign equal numbers of participants to each group and actively minimise the variance of MVC values across the protocols.

#### Assessment task (AT)

To assess the effect of different force intensities of unimanual fatigue on the movement quality of both arms, participants completed a bimanual object-hit task for a duration of 4min. The task was adapted and distinctly modified from the original object-hit task by Tyryshkin et al^17^. To complete the task, participants needed to perform arm reaches with robotic handles (displayed to the participant as virtual green paddles of 1.5 cm width and 0.5 cm height) to deflect virtual red balls (0.75 cm diameter) away (**Figure 1b**). Balls were continuously released from 16 evenly distributed bins at the rear/top side of the display (30 objects per bin), falling toward the participant’s side of the display at different velocities (15-30 cm/s) with two balls per second. The bin and speed of the falling virtual objects were kept continuously pseudorandom and unpredictable to the participant. The comparably small width of the paddles was chosen to increase the accuracy of movements required to hit a ball. When the participant hit a ball with the paddle, the direction of the ball movement changed, providing real-time feedback to the participant.

#### Recovery period

The recovery period between task rounds was 20 minutes, which has been shown to be adequate to allow for good restitution of most elements of voluntary muscle force activation^18^. During the first recovery period after the first experimental round, participants were shown a standardised 14 min section from a documentary movie (‘Planet Earth’, S01). During the second recovery period after the second experimental round, participants then watched the subsequent 14 min section from the same movie.

### 2.3. Data acquisition

#### Force data acquisition

MVC and FT were conducted using a hand dynamometer (Biopac Systems Inc., Aero Gamino Goleta, CA, USA) and custom code implemented in *MATLAB R2022a* (The MathWorks Inc., Natick, MA, USA) using the Psychophysics Toolbox extensions (*Psychtoolbox-3)*^19,20^. Force data were sampled at 47.75 Hz. Dynamometer force data were routed through Lab Streaming Layer (LSL) and LabRecorder (https://github.com/labstreaminglayer/App-LabRecorder).

#### Kinematic data acquisition

Kinematic movement quality assessments were acquired using a *Kinarm End-Point Lab* robot with *Dexterit-E 3.9.2* software (BKIN Technologies Ltd., Kingston, ON, Canada).

#### EMG data acquisition

During performance of MVC, FT, and AT, surface electromyography (EMG) of the right and left flexor carpi radialis (FCR) and extensor carpi radialis (ECR) muscles was acquired using bipolar configuration with a TMSi Porti7 recorder (Twente Medical Systems International B.V., Oldenzaal, The Netherlands). Small adhesive disposable electrodes with a contact area of 4.5cm^2^ (Kendall H124SG; Cardinal Health, Dublin, OH, USA) were used, and placed in a bipolar setup with two electrodes placed next to each other on the bulk of the muscle belly at a fixed distance of 24mm between the centres of the electrodes. Participants were instructed to perform a handgrip contraction during which the muscles were palpated manually to determine correct placement. A common grounding electrode was placed on the ulnar styloid process. EMG signals were recorded in raw format at a sampling rate of 2048 Hz and were routed through LSL and LabRecorder.

### 2.4. Data analysis

#### MVC data

Median force was calculated for each MVC separately, by calculating the median force applied between the full width at half maximum (FWHM) points of each contraction force profile. (**Figure 1d**). For MVC 3× the highest of the 3 median values of the 3 MVCs from each participant was used for statistical analysis. The highest median value of the MVC 3× at baseline was used as a reference value for computing subsequent target values for the FT.

#### Force data

The first 10 of the 270 separate contractions of each FT were classed as a familiarisation period and hence excluded from the analysis. For the remaining 260 contractions the area under the curve (AUC) was computed over the 1 sec contraction window, as shown before.^3^ Next, the slope of the 260 AUC values was computed using linear regression. The derived beta value of the regression was subsequently utilised for force output quantification, with a negative beta value indicating declining force output, which represents one of the main criteria for identifying muscle fatigue.

#### Kinarm data

From the kinematic data, the number of hits per hand was calculated as the primary outcome measure. We also calculated a number of secondary outcome measures, including velocity, (absolute) acceleration, and space covered for both hands separately. Space covered was defined as the area a hand covered across the entire task duration and calculated for the left and right hand separately.

#### EMG data

Using MATLAB R2022a (The MathWorks Inc., Natick, MA, USA) the EMG data were bandpass filtered between 10 Hz and 500 Hz with a fourth-order Butterworth filter, and the envelope of the EMG data was computed. During the FT the EMG data were analysed in a similar manner to the Force data, i.e., we removed the first 10 contractions, calculated the AUC per contraction, and then computed a linear regression across contractions. During the AT we computed the median of the envelope from the beginning to the end of the task.

### 2.5. Statistical analysis

Statistical analyses were performed using Prism version 10.2.2 (Graphpad), JASP version 0.18.3 (University of Amsterdam, The Netherlands) and MATLAB *R2022a* (The MathWorks Inc., Natick, MA, USA). Differences between the three different force levels of the FT (5%, 50%, 75%) on FT and AT performance were investigated using repeated-measures Analysis of Variance (RM-ANOVA) and follow-up t-tests. We tested whether the force level of the FT affected the space covered by the right (fatigued) or left (unfatigued) hand via a two-samples t-test using cluster-based non-parametric permutations with row-shuffle permutations (5000 permutations)^21^. A cluster forming threshold of p = 0.01 was used and a spatial extent threshold of 1 was set to ensure that any cluster has at least 2 adjacent points exceeding the cluster forming threshold. Partial eta-squared (ηp^2^) and Cohen’s d (*d*) effect sizes are reported for RM-ANOVA and t-tests respectively. Data visualisations were generated using Prism version 10.2.2 (Graphpad), MATLAB *R2022a* (The MathWorks Inc., Natick, MA, USA), Adobe Illustrator CC 2024 and Adobe Photoshop CC 2024 (Adobe Inc, San Jose, CA, USA).

## 3. Results

### 3.1. Force levels during the Fatiguing Task affect several behavioural and physiological outcomes

To test whether the three different FT force levels (5%, 50%, 75% MVC) had the expected differential effects on behavioural and physiological markers of fatigue, we first needed to test whether the MVCs acquired before the FT were comparable across the three rounds. A 1 × 3 RM-ANOVA with a within-subjects factor force level (5%, 50%, 75% MVC) demonstrated no significant main effect of force level on MVC (*F*_2,58_ = 0.174, *p* = 0.841, η_p_^2^ = 0.006), suggesting that the MVC did not vary between conditions (5% MVC: *M* = 57.233, *SD* = 17.938; 50% MVC: *M* = 58.178, *SD* = 18.039; 75%: *M* = 57.060, *SD* = 20.041). To test whether the order in which participants performed the three different FT force levels impacted on their FT performance, we added a between-subjects effect of order, and saw no significant main effect (*F*_5,24_ = 0.414, *p* = 0.83, η_p_^2^ = 0.079) and no order × force interaction (*F*_10,48_ = 1.68, *p* = 0.113, η_p_^2^ =0.259).

#### Performance on the Fatigue Task is inversely related to force level

We first wanted to test whether different force levels had different effects on FT performance, as quantified as the slope of the linear regression across the 260 contractions (i.e., AUC values). We tested whether the FT performance was affected by force-level using a 1 × 3 RM-ANOVA with a within-subjects factor of force level (5%, 50%, 75% MVC) which demonstrated a significant main effect of force level (5% MVC: *M* = -0.023, *SD* = 0.229; 50% MVC: *M* = 1.305, *SD* = 1.305; 75%: *M* = -2.363, *SD* = 2.208; *F*2,58 = 22.747, *p* < 0.001, η_p_^2^ = 0.440, **Figure 2a**). All follow-up *t*-tests were significant (5% vs 50% MVC: *t*_29_ = 3.427, *p*_bonf_ = 0.003, *d* = 0.800; 5% vs 75% MVC: *t*_29_ = 6.745, *p*_bonf_ < 0.001, *d* = 1.57; 50% vs 75% MVC: *t*_29_ = 3.318, *p*_bonf_ = 0.005, *d* = 0.77), suggesting that, as expected, the participant’s FT performance deteriorated with increasing force level.

**Figure 2:**
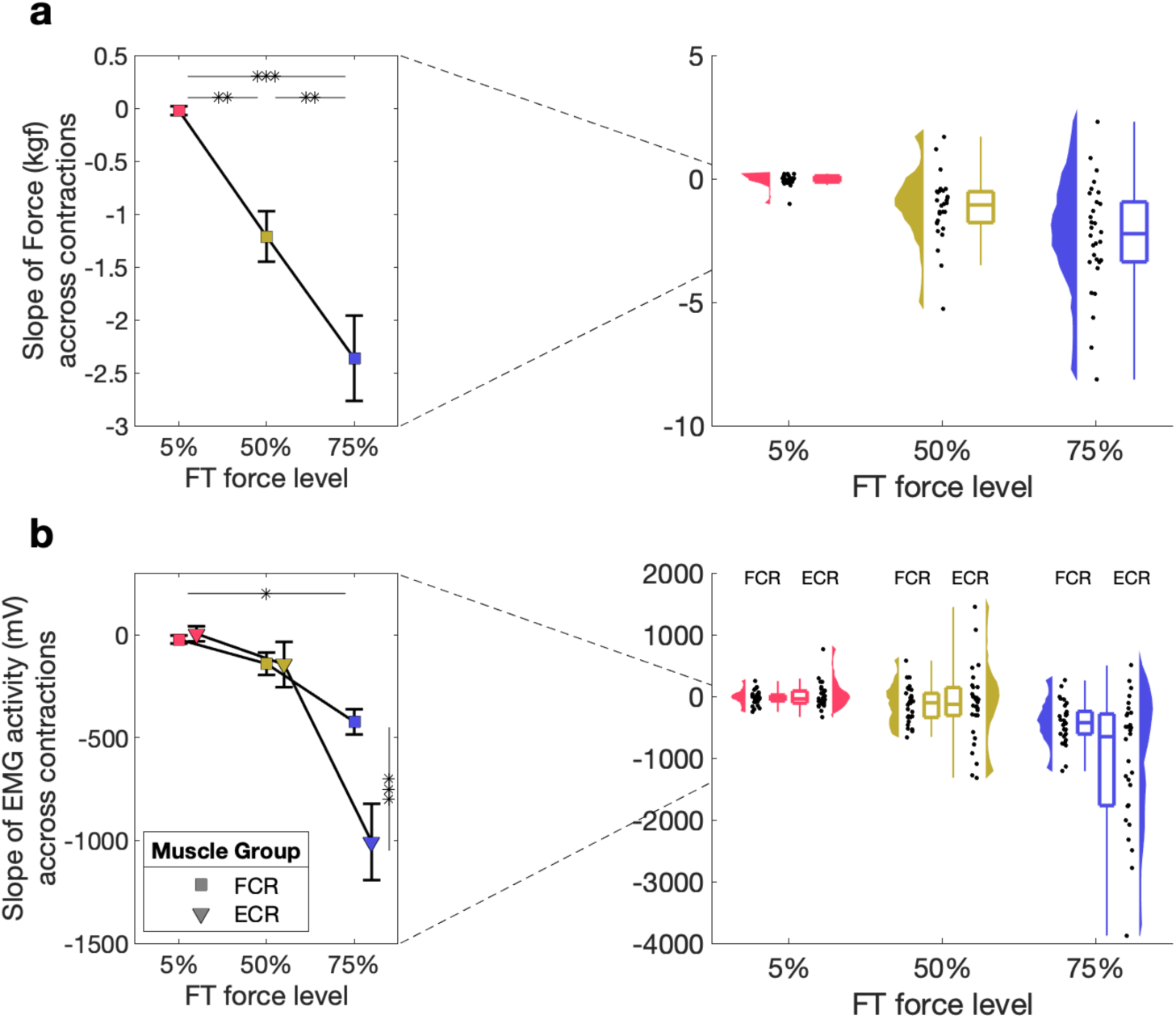
Direct measures of fatigue during the FT. a) FT performance for 5%, 50%, 75%. Mean and standard error across subjects is shown (left) as well as single subject data, distribution, and boxplot (right). b) FT muscle activity from the FCR and ECR for 5%, 50%, 75%. Mean and standard error across subjects is shown (left) as well as single subject data, distribution, and boxplot (right).

#### Decrease in EMG activity during the Fatiguing Task only occurs at 75% MVC

To investigate the effects of our FT on muscle activity, we also evaluated EMG activity from two muscles during FT using a 2 × 3 RM-ANOVA with within-subject factors of muscle (FCR, ECR) and force level (5%, 50%, 75% MVC; **Figure 2b**). This revealed a significant main effect of muscle (*F*_1,59_ = 8.658, *p* = 0.006, η_p_^2^ = 0.230) and force level (*F*_2,58_ = 24.263, *p* < 0.001, η_p_^2^ = 0.456), as well as a significant muscle × force level interaction (*F*_2,58_ = 13.190, *p* < 0.001, η_p_^2^ = 0.313). Follow-up tests suggest that these differences are driven by significant differences at 75% MVC, suggesting that the magnitude of EMG activity only decreases during with the highest FT.

#### MVC is decreased after both 50% and 75% MVC force levels

Next, we examined how the ability to produce maximal contractions is affected by the FT, as quantified by the MVC performed directly after the FT. A 1 × 3 RM-ANOVA with a within-subjects factor force level (5%, 50%, 75% MVC), demonstrated a significant main effect of force level (5% MVC: *M* = 52.094, *SD* = 14.771; 50% MVC: *M* = 41.142, *SD* = 15.134; 75%: *M* = 41.395, *SD* = 17.721; *F*2,58 = 13.892, *p* < 0.001, η_p_^2^ = 0.324; **Figure 3a)**. Follow-up *t*-tests revealed significant differences between 5% and 50% MVC (*t*29 = 4.617, *p*_bonf_ < 0.001, *d* = 0.69) as well as between 5% and 75% MVC (*t*29 = 4.511, *p*_bonf_ < 0.001, *d* = 0.67), but not between 50% and 75% MVC (*p*_bonf_ > 0.1), suggesting that 50% and 75% force levels similarly effect the MVC.

**Figure 3:**
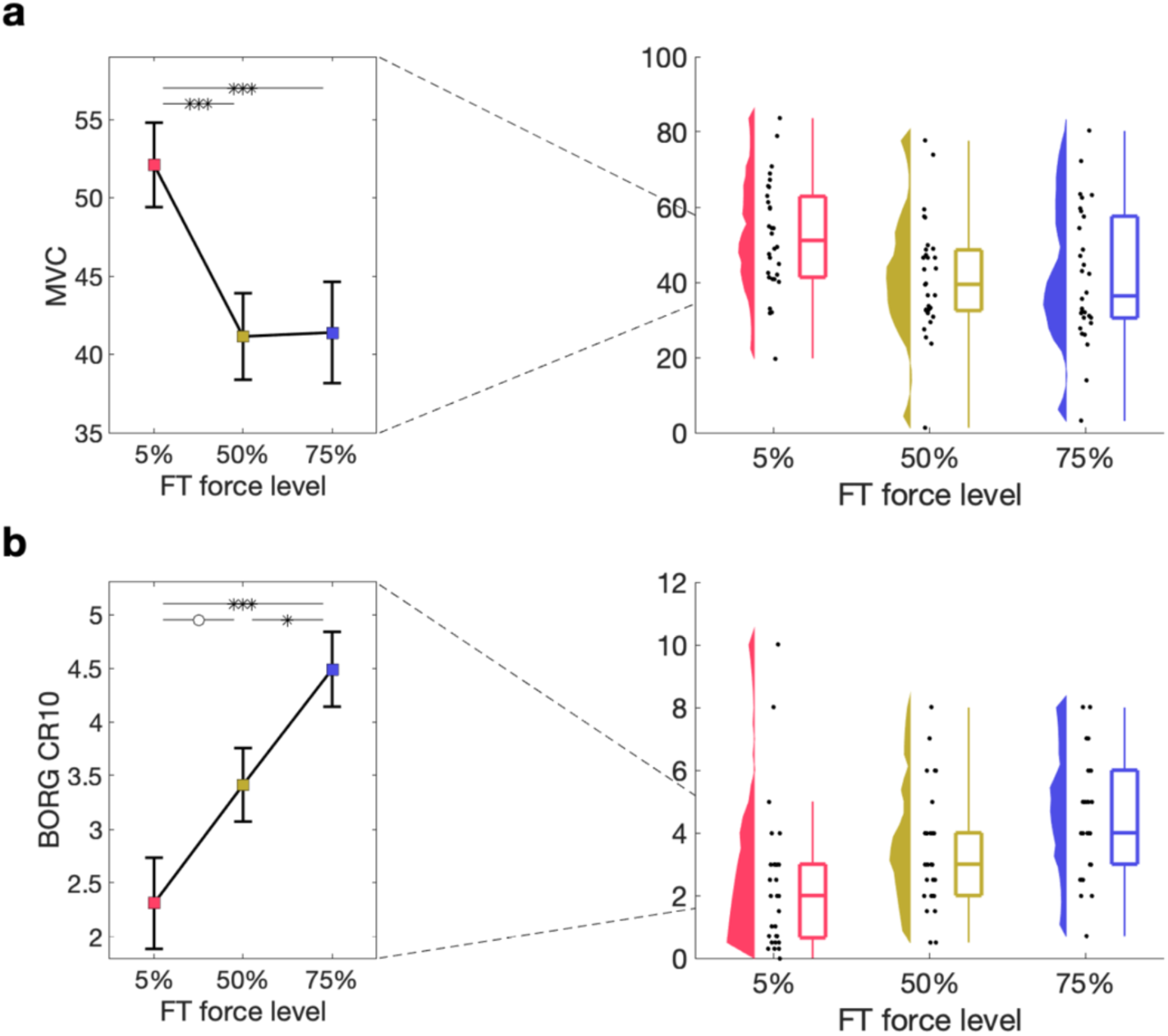
Indirect measures of fatigue during the FT. a) MVC directly after the FT for 5%, 50%, 75%. Mean and standard error across subjects is shown (left) as well as single subject data, distribution, and boxplot (right). b) Perceived exertion quantified using the BORG CR10 after the FT for 5%, 50%, 75%. Mean and standard error across subjects is shown (left) as well as single subject data, distribution, and boxplot (right).

#### Perceived exertion increases with force level of FT

Lastly, we examined how perceived exertion, as indexed by the BORG CR10 obtained at baseline and after the FT, was affected by different force levels during the FT. A 1 × 3 RM-ANOVA with a within-subjects factor of force level (5%, 50%, 75% MVC) showed a significant main effect of force level on BORG CR10 (5% MVC: *M* = 2.567, *SD* = 2.650; 50% MVC: *M* = 3.500, *SD* = 1.871; 75%: *M* = 4.507, *SD* = 1.847; *F*2,58 = 12.787, *p* < 0.001, η_p_^2^ = 0.306; **Figure 3b**). Follow-up *t*-tests revealed significant differences between 5% and 75% MVC (*t*29 = -5.056, *p*_bonf_ < 0.001, *d* = -0.90) as well as between 50% and 75% MVC (*t*29 = -2.623, *p*_bonf_ = 0.033, *d* = -0.47) with a trend between 5% and 50% MVC (*t*29 = -2.432, *p*_bonf_ = 0.054, *d* = -0.43), suggesting that perceived exertion increases with increasing FT force level.

### 3.2. Force levels during the Fatigue Task affect the magnitude of EMG activity, but not kinematics, during the Assessment Task

Next, we examined whether the three different force levels (5%, 50%, 75% MVC) of the FT affected performance and muscle activity during AT for both the right (fatigued) and left (unfatigued) hands separately.

#### No significant effect of fatigue on number of targets hit

Figure 4a shows the hits per bin during the AT for the three FT force levels. A 2 × 3 RM-ANOVA with a within-subject factor of hand (right [fatigued], left [unfatigued]) and force level (5%, 50%, 75% MVC) demonstrated no significant main effect of hand or force level, and no significant hand × force level interaction (all *p’s* > 0.1).

**Figure 4:**
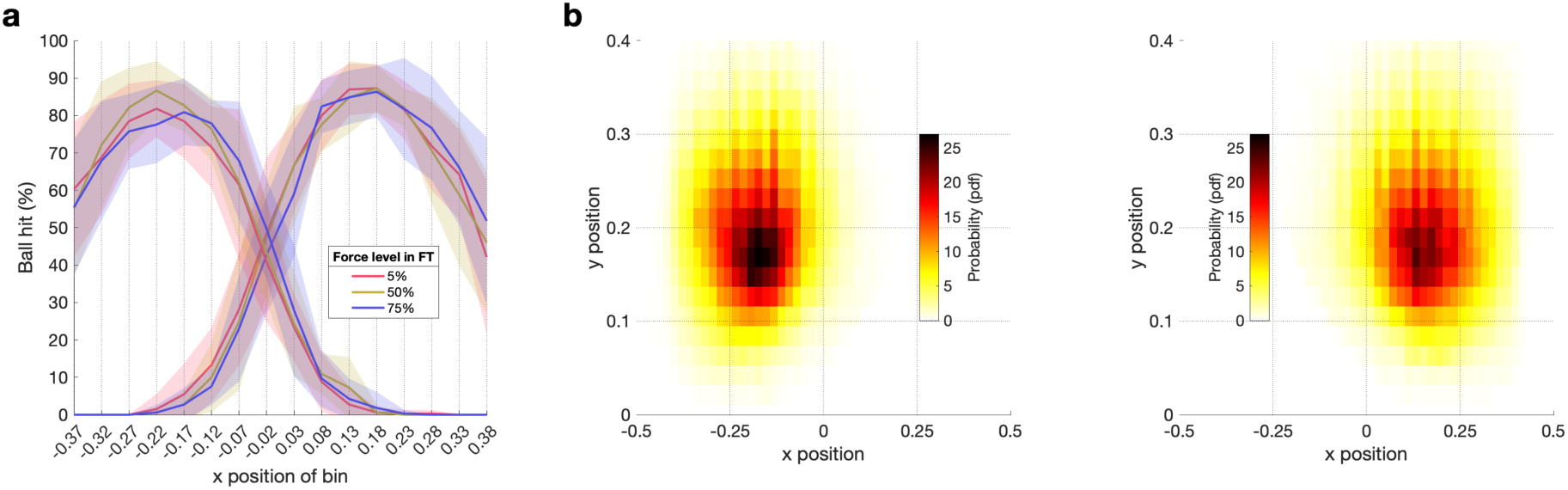
Ball hits and space covered per hand during the AT. a) Ball hits per bin for the left (unfatigued) hand and the right (fatigued) hand during the AT performed after the FT for 5%, 50%, 75%. The solid line represents the mean across subjects, shaded area represents the standard error across subjects. b) Heat map of the space covered by the left (unfatigued) hand and the right (fatigued) hand during the AT. Data are averaged across subjects and force levels of FT for 5%, 50%, 75%.

#### No significant effect of fatigue on space covered

Next, we examined how the space covered during the AT by the right (fatigued) and left (unfatigued) hand is affected by different force levels during the preceding FT. Figure 4b shows the grand average heatmaps per hand. There were no significant differences between conditions for either hand.

#### No significant effect of fatigue on acceleration and velocity in AT

We then assessed if acceleration and velocity of the right (fatigued) and left (unfatigued) hand were affected by different FT force levels, using two separate 2 × 3 RM-ANOVA tests with the within-subject factors of hand (right [fatigued], left [unfatigued]) and force level (5%, 50%, 75% MVC). We found no significant main effect of hand or force level, and no significant hand × force level interaction on velocity (all *p’s* > 0.1, Figure 5a). In terms of acceleration, we found a significant main effect of hand with higher acceleration for the left hand (*F*_1,29_ = 433.389, *p* < 0.001, η_p_^2^ = 0.881), but no significant main effect of force level (*p* > 0.1), and no significant hand × force level interaction (*p* > 0.1, Figure 5b).

**Figure 5:**
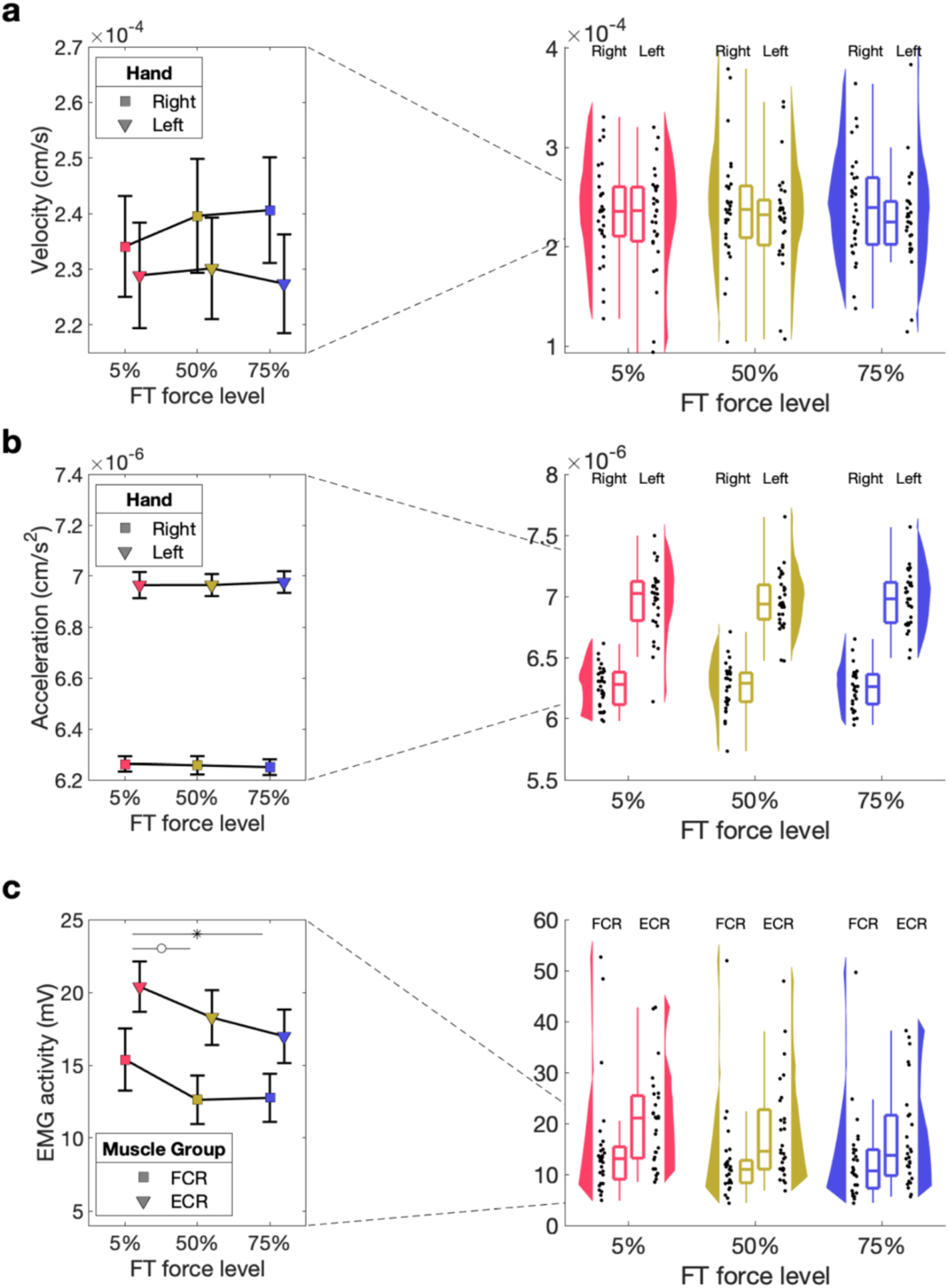
Kinematic measures and EMG activity during the AT. a) Velocity of the fatigued (right) hand during the AT performed after the FT for 5%, 50%, 75%. Mean and standard error across subjects is shown (left) as well as single subject data, distribution, and boxplot (right). b) Acceleration of the fatigued (right) hand during the AT performed after the FT for 5%, 50%, 75%. Mean and standard error across subjects is shown (left) as well as single subject data, distribution, and boxplot (right). c) EMG activity of the fatigued (right) hand during the AT performed after the FT for 5%, 50%, 75%. Mean and standard error across subjects is shown (left) as well as single subject data, distribution, and boxplot (right).

#### Assessment Task – Magnitude of EMG activity

Finally, we wanted to test whether our fatiguing task made a significant difference to muscle activity in the AT, in the absence of behavioural effects. We therefore performed 2 × 3 RM-

ANOVA tests with the within-subject factor of muscle (FCR, ECR) and force level (5%, 50%, 75% MVC) for each hand separately. For the right (fatigued hand) this showed a significant main effect of force level (*F*_2,54_ = 4.982, *p* = 0.010, η_p_^2^ = 0.156), a trend for the main effect muscle (*F*_1,27_ = 4.090, *p* = 0.053, η_p_^2^ = 0.132), but no significant muscle × force level interaction (*p* > 0.1; Figure 5c). There was no significant main effect of force level or a significant muscle × force level interaction (both *p’s* > 0.1) for the left (unfatigued) hand.

## 4. Discussion

In this study, we investigated the effects of unilateral hand grip fatigue on the movement quality of both arms in a bilateral task. To that end, we utilised a variety of methods to probe

1. whether the FT protocol we employed adequately elicited unilateral muscle fatigue, thereby creating the desired experimental precondition for testing our main hypotheses and
2. how the achieved unilateral muscle fatigue then affected a range of movement quality measures in a bilateral movement task (AT).

### 4.1. Our fatiguing task induced fatigue across multiple outcome measures

To establish whether our FT did indeed induce fatigue in the target muscles (FCR and ECR) we used a combination of metrics each of which are sensitive to detecting fatigue. We showed that the higher force levels for our FT, which were designed to induce fatigue, did indeed induce a decline in real-time force output during the task^22^, characteristic changes in the magnitude of EMG activity during the task, with some increases in magnitude seen at lower force intensities and progressive magnitude decline at higher force intensities^23^, a decline in post-exercise MVC^1^, and an increase in the self-reported perceived rate of exertion^24^. Collectively, significant changes were observed across all four metrics with increasing force levels, indicating that muscle fatigue occurred with the FT protocol.

In addition, while force performance data showed significant differences between all force levels, EMG activity did not significantly differ between 5% MVC and 50% MVC force level, only decreasing significantly at 75% MVC. This may reflect a physiological attempt at compensation in response to emergent handgrip fatigue at a moderate force level, a mechanism likely mediated by increased muscle fibre recruitment^23^. In the 75% MVC condition, this compensation then seemed to fail most drastically in the ECR, when compared to the FCR. Interestingly, when comparing post-exercise MVC measures, there is a comparable decline from control for both 50% and 75% MVC force levels with no significant difference between the two fatigue conditions. When contrasting this finding with the real-time force performance data, this might indicate that higher force level exercise seems to affect repeated submaximal contraction performance more dramatically than maximum post-exercise contraction performance. This finding might be in part driven by motivation, as participants might have felt a greater sense of achievement or relief after having completed the 75% MVC task versus after the 50% MVC task, possibly influencing their subsequent resolve to make maximal contraction efforts on the post-FT MVC test. The self-reported measures, on the other hand, showed a more linear trend of increasing perceived level of exertion with increasing force level.

In summary, these findings indicate that in our experimental setting, muscle fatigue was achieved in both experimental conditions compared to the control condition with some degree of scaling between 50% and 75% of MVC detectable in three of the four available measures (force output, magnitude of EMG activity, and subjective exertion).

### 4.2. Handgrip FT elicits sustained ipsilateral electromyographical effect throughout AT

Through analysis of simultaneously recorded EMG signal magnitude from the right arm during the AT, we conclude that the FT at both experimental force levels (50% and 75% of MVC) results in an ipsilateral reduction of EMG activity on the subsequent AT compared to the control condition (5% of MVC). This indicates that the implemented fatigue protocol shows a sustained fatiguing effect throughout the AT. Interestingly, the difference between the ECR and the FCR muscle seen during the FT at 75% MVC force level appears to largely disappear in the subsequent AT. One must consider, however, that 1) the continuous method of measuring EMG activity on AT in contrast to the discrete, contraction-based EMG measurements on FT might render these outcomes difficult to compare and 2) the comparison of EMG activity between FT and AT is a non-specific comparison given the movement type differences in the two tasks.

### 4.3. Ipsilateral movement quality in an object-hit task remains unimpeded by dominant handgrip fatigue

Even though participants continued to show signs of fatigue in their right arm as quantified by EMG during the AT, this did not significantly influence hits, space covered, velocity or acceleration of the fatigued (ipsilateral) arm. Therefore, contrary to our expectations, it can be concluded that sustained unilateral handgrip fatigue does not impair ipsilateral arm movement quality in a bilateral object-hit task performed on the Kinarm endpoint robot. One possible explanation for this finding is that the employed handgrip FT is not kinematically specific to the following AT. Although handgrip exercises tend to elicit activity in most muscles of the upper extremity, the proportion of muscle involvement is different when compared to the muscles involved in the movements necessary to hit balls on an object-hit task. The lack of performance deterioration of the fatigued ipsilateral (right) arm despite handgrip fatigue could, in turn, also explain why no significant spillover of activity from the left arm to the right area of the canvas could be observed - there was simply no need for the unfatigued (left) arm to support the tasking of the fatigued homologue.

### 4.4. Contralateral higher-level movement quality is unchanged following dominant handgrip fatigue

Based on past experiments by our group showing that unilateral handgrip contractions at 50% of MVC were associated with improved motor response latency in the contraleral, unfatigued homologue and related changes in cortical GABA-ergic activity, we aimed to test whether these effects translate into measurable effects in higher-level movement quality measures. Neither the primary outcome (number of balls hit), nor secondary outcomes (space covered, velocity and acceleration during object-hit task) showed any significant change on the side of the contralateral, unfatigued arm. Therefore, we can conclude from the data presented in this study that there is no measurable effect of unilateral handgrip fatigue on higher-level movement quality on the contralateral homologue. The significant effects seen in our previously employed button-press task^3^ did not translate into advanced movement quality benefits.

### 4.5 Clinical implications and outlook

Fatigue is very common in patients with pathological unilateral motor impairment, e.g., after surviving a stroke; indeed it is among the most regularly recorded symptoms of stroke survivors.^25^ Although post-stroke fatigue is currently believed to be caused mainly by impaired corticomotor excitability,^25^ rendering it mechanistically different from physiological muscle fatigue, some characteristics of post-stroke fatigue overlap with physiological fatigue. In post-stroke fatigue, a decrease in corticomotor neural drive after surviving a stroke creates a scenario of disuse or limited use of the affected muscles, leading to atrophy (especially with regards to type II muscle fibres), diminished ability to voluntarily contract muscles and less synchronization of motor units^26^, effects which, to a certain degree, can also be observed in healthy people, e.g. after physical inactivity^27^. In any case, gaining a better understanding of how neural dynamics shape the adaption response to unilateral fatigue is not only of fundamental scientific interest but also of clinical relevance.

Interestingly, while our findings did not demonstrate a measurable improvement in high-level movement quality in the contralateral limb, the lack of negative impact on performance in the unfatigued arm could have positive clinical implications. For instance, our results suggest that unilateral handgrip fatigue training does not impair contralateral movement quality, highlighting a potential strategy for strength training and neuromodulatory benefits in rehabilitation settings. Research has shown that high-intensity interval exercise can promote systemic physiological benefits, including increases in brain-derived neurotrophic factor (BDNF)^28^ and other neurotrophic factors that support neuroplasticity and functional recovery. While it needs to be noted that our study was conducted in healthy volunteers and only measured movement quality on the same day as the fatiguing task intervention, the finding of a lack of negative impact still opens up the possibility of incorporating handgrip exercises or similar unilateral fatigue-inducing tasks on the unimpaired limb before or alongside targeted rehabilitation exercises for the impaired limb. By doing so, patients might gain neurophysiological benefits that enhance the neural state without compromising the quality of rehabilitation in the affected limb. This point highlights the potential clinical efficacy of ‘cross-education’, whereby single limb strength and/or skilled motor training confers a positive training affect to the contralateral homologous muscle group.^29^ This approach could potentially maximize the benefits of the paretic limb,^30^ but also boost systemic neuroplastic adaptations, setting a stronger foundation for motor recovery in the affected limb, while leaving high-level motor function intact in the unfatigued, contralateral limb. Further studies

could explore this hypothesis, examining how such protocols might be refined and applied in clinical rehabilitation settings to support motor recovery and neuroplastic gains.

## Acknowledgments

We thank Prof. Huiling Tan and Dr. Alek Pogosyan for sharing their EMG equipment.

## Funding

The study was supported by a Senior Research Fellowship awarded to CJS, funded by the Wellcome Trust (224430/Z/21/Z). ALK was funded by a Medical Research Travel Grant from the Harold Hyam Wingate Foundation. JWA is supported by an internally funded Vice-Chancellor Fellowship at Northumbria University. SS was supported by the Jon Moulton Charity Trust. This work was part funded by the MRC Brain Network Dynamics Unit at the University of Oxford (MC_UU_00003). The research was supported by the National Institute for Health Research (NIHR) Oxford Biomedical Research Centre. The work was supported by the NIHR Oxford Health Biomedical Research Centre (NIHR203316). The views expressed are those of the author(s) and not necessarily those of the NIHR or the Department of Health and Social Care. The Wellcome Centre for Integrative Neuroimaging is supported by core funding from the Wellcome Trust (203139/Z/16/Z and 203139/A/16/Z). For the purpose of open access, the author has applied a CC BY public copyright licence to any Author Accepted Manuscript version arising from this submission.

## Author contributions

ALK: Conceptualization, Investigation, Formal Analysis, Project Administration, Visualization, Writing – Original Draft Preparation

JWA: Conceptualization, Writing – Original Draft Preparation

SS: Conceptualization, Methodology, Writing – Review & Editing

CJS: Funding Acquisition, Resources, Writing – Review & Editing

CZ: Conceptualization, Data Curation, Formal Analysis, Project Administration, Resources, Visualization, Writing – Original Draft Preparation

## Competing interests

The authors report no competing interests.

## Notes

### Competing Interest Statement

The authors have declared no competing interest.

